# Identification of bacteria involved in the decomposition of lignocellulosic biomass treated with cow rumen fluid by metagenomic analysis

**DOI:** 10.1101/570374

**Authors:** Chol Gyu Lee, Yasunori Baba, Ryoki Asano, Yasuhiro Fukuda, Chika Tada, Yutaka Nakai

## Abstract

Previously, pretreatment of plant biomass using rumen fluid systems was developed to decompose cell wall. However, microbes which involved in plant cell wall decomposition in this system have not been identified, because the conditions of this system are different from the *in situ* rumen environment. We investigated the bacteria involved in the decomposition of cellulose and hemicellulose in a waste paper with the rumen pretreatment system using shotgun metagenomic analysis with next generation sequencing. After pretreatment of waste paper, about a half of the cellulose and hemicellulose content was decomposed. Genes encoding for cellulase and hemicellulase were mainly found to belonging to *Ruminococcus, Clostridium*, and *Exiguobacterium*. This study shows that *Clostridium* and *Exiguobacterium*, which have not been identified as predominant genus involved in cellulose and hemicellulose decomposition, might be categorized as the main fibrolytic bacteria in this system.

## Introduction

The development of carbon-neutral biomass conversion processes has attracted interest in recent years, promising an alternative platform for the production of fuels and chemicals that could reduce the global dependence on petroleum [7]. Lignocellulosic plant biomass is the most abundant carbon source on the earth has potential to be a renewable carbon [27]. The plant cell wall is composed of three closely related polymers: cellulose, hemicellulose, and lignin. They form a physical barrier that inhibits the adsorption of hydrolytic enzymes and protects them. For effective carbon utilization, plant biomass has been pretreated with wet oxidation and chemical methods so the plant cell wall can be hydrolyzed to monosaccharide [14, 38]. However, wet oxidization requires a substantial amount of energy and chemical methods developed so far are inefficient for lignocellulose decomposition [9, 21].

Ruminant animals efficiently utilize plant materials as an energy source because they can decompose plant cell walls in the rumen. Microorganisms in the rumen play an important role in hydrolysis and fermentation of lignocellulose [19, 20]. Several studies have indicated that rumen microbes can degrade a larger amount of cellulose in less time than anaerobic digester [25, 31]. Baba et al. [2] developed an effective lignocellulosic biomass decomposition method that uses rumen fluid. It can decompose more than half of the cellulose, hemicellulose, and lignin content by pretreating with rumen fluid over 24 h. Baba et al. [1] further discovered that the bacterial community changes during the pretreatment period by 16S rRNA amplicon sequencing analysis. While, this analysis has been clarified for the microbial composition, but it did not explicitly identify the microorganisms involved in the decomposition process. Shotgun metagenomic analysis randomly sequencies the DNAs present in environmental samples using a next-generation sequencer, which provides an effective method to determine microbial composition and general function from uncultivated microorganisms [33]. Several studies have conducted metagenomic analysis in cow rumen and revealed the several microbes that are involved in the decomposition of the plant cell wall decomposition [3, 5]. However, the pretreatment process is an unbuffered closed system that treats a single substrate. Further complicating the system, different microbes may be involved in the *in vitro* decomposition process compared with *in situ* cow rumen environment. Thus, we conducted a shotgun metagenomic study on the lignocellulosic biomass pretreatment with rumen fluid system to reveal the cellulose and hemicellulose decomposing microbes. This study shows that unique this system may utilize unique microbes compared to the *in situ* rumen environment to decompose the plant cell wall in waste paper.

## Material and Method

### Pretreatment of wastepaper using rumen fluid

The treatment condition followed the method of Baba et al. [2]. Rumen contents were collected orally from grass-fed cattle and filtered with a mesh strainer (1 mm × 1 mm) to remove coarse solids. 0.3g of the waste paper pieces (35 mm long × 25 mm wide) were soaked in 300 ml of rumen fluid containing 0.3 g of L-cysteine as a reducing agent. They were incubated at 37 °C on a rotary shaker at 140 rpm for 120 h. Fluid samples of the reaction were collected before and after 6, 24, 48, 60, 96 and 120 h of pretreatment and conducted to chemical analysis. The experiments were conducted in duplicate.

### Chemical composition of the waste paper and pretreatment systems

The cellulose, hemicellulose, and lignin contents in the waste paper were determined according to the detergent fiber method [34]. The chemical composition of the waste paper was 71.3% cellulose, 18.5% hemicellulose, and 8.9% lignin. Volatile fatty acid (VFA) concentrations were determined using LC-10Avp Tvp HPLC with an ultraviolet detector (Shimadzu, Kyoto, Japan), equipped with an ion-exchanger column RSpak KC-811 (Shodex, Tokyo, Japan). Analysis were performed at 60 °C using 3 mM HClO_4_ for elion. The dissolved chemical oxygen demand (dCOD) was measured by the dichromate reactor digestion method [15].

### DNA extraction from rumen fluid

The fluid samples collected from before and after 60 h pretreatment were used for DNA extraction because microbes actively decompose substrates in this time point (see results). DNA was extracted from microcosms according to the method of Morita et al. [23]. In brief, bacterial DNA was isolated by the enzymatic lysis method. Cells were lysed in the presence of lysozyme, achromopeptidase, and Proteinase K. Cleared lysate was treated with phenol/chloroform/isomyl alcohol and RNase. DNA was precipitated with ethanol and suspended dissolved in Tris-EDTA (TE) buffer. DNA was quantified using PicoGreen™ dsDNA Assay Kit (Thermo Scientific Massachusetts, USA). The integrity of DNA was verified by a Nanodrop (Thermo Scientific, Massachusetts, USA) and gel visualization (0.8 % agarose in Tris-acetate EDTA buffer).

### Shotgun metagenome sequencing

DNA Libraries were subjected to paired-end sequencing on an Illumina MiSeq sequencer (Illumina, SanDiego, CA, USA) at the Biotechnology Center, Akita Prefectural University (Akita, Japan) with Miseq Reagent Kit v2 500 cycles (∼ 250 bp) using Truseq library kit (Illumina) according to the manufacturer’s instructions. The information obtained by sequencing is shown in Table S1. The sequencing data were deposited in the DNA Data Bank of Japan (DDBJ) Sequence Read Archive under accession number is DRA007782.

### Data analysis

Paired-end sequences were joined together with > 30 bp overlapping sequences containing < 10% mismatches and then converted from fastq to the fasta format with a Q-score cutoff of > 30 using PRINSEQ [29]. Ribosomal RNA sequences (rRNAseq) in the metagenomics data were identified using a BLAST search against the M5RNA database (E value of < 10^−5^, alignment length of >50 bp) on the MG-RAST server ver. 3.6 [22]. Microbial diversity and richness were shown as Shannon index, operation taxonomic units (OTU) numbers, and Chao1 as index determined by MG-RAST. We collected highly abundant gene sequences of key enzymes involved in cell wall decomposition according to the KEGG compound database [16]. The identified sequences were compared to the NCBI nr database in December 2015 by BLASTX for more precise assignment (E value of < 10^−5^, alignment length of >30 aa).

## Results

### Pretreatment of lignocellulosic biomass using rumen fluid

Approximately, half of the cellulose, hemicellulose, and lignin in waste paper reactions were decomposed after 60 h of pretreatment using rumen fluid (Table 1). The decomposition increased dCOD and two of the three majored VFAs, which correlated with a decreasing pH value (Figure 1) similar to Baba et al. [1, 2] using waste paper and rapeseed as substrates. VFA concentrations and dCOD were continuously increased during pretreatment until 120 h. These results indicated that microbes were actively decomposing the waste paper at the 60 h time point.

**Table 1.**
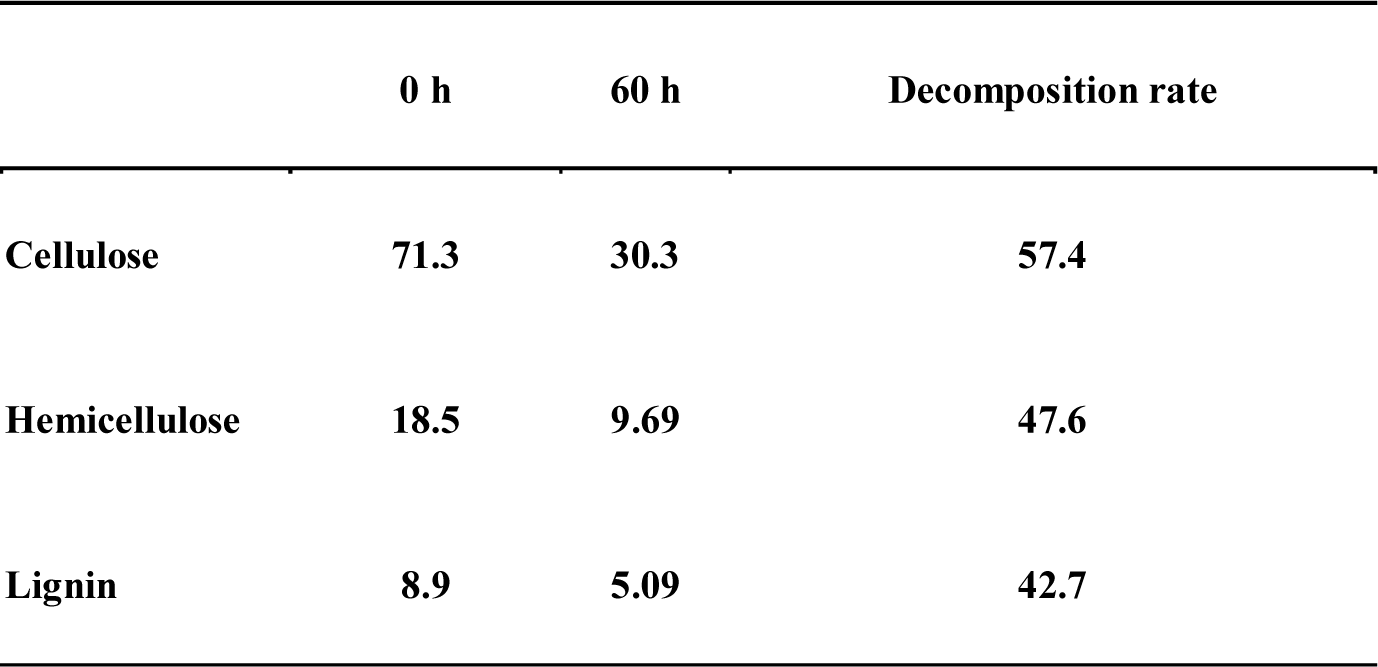
Chemical components (%) of waste paper before and after pretreatment

**Figure 1.**
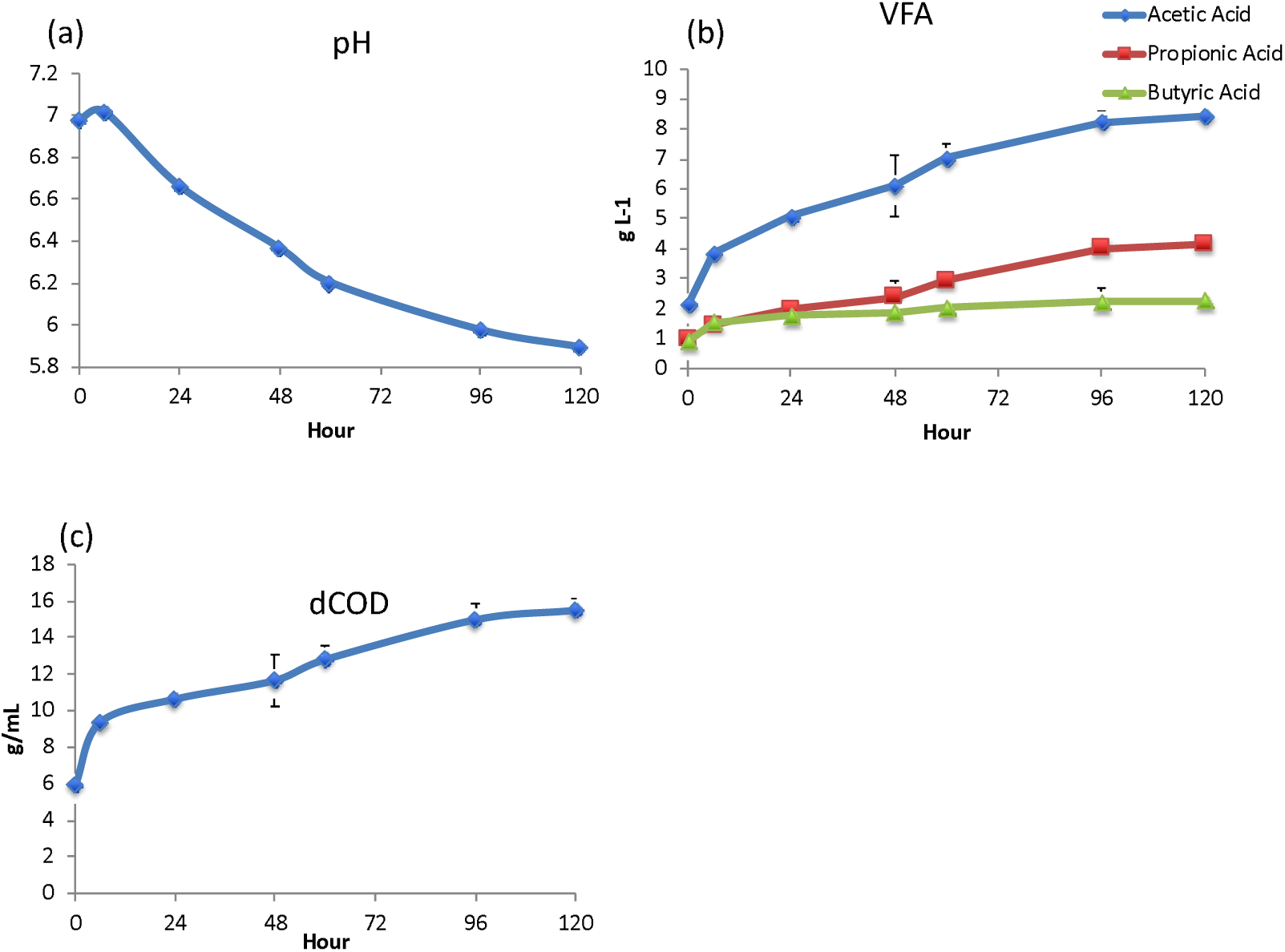
Chemical properties of the rumen fluid system during the incubation period. (a) pH; (b) Volatile fatty acids concentration; (c) Dissolved chemical oxygen demand (dCOD) concentration

### Alteration of microbial community, diversity, and richness in the pretreatment system

The microbial communities residing in the rumen fluid changed during the incubation period (Figure 2). At the phylum level, Firmicutes were increased from 53.4 and 65.5 % before information to 66.5 and 75.0 % after incubation in test sample one and two, respectively. At the class level *Clostridia*, a class of Firmicutes represented 36.4 and 50.3 % before incubation to 50.3 and 63.6 % after incubation. The relative abundance of other classes of microbes, including Bacilli, Bacteroida and Gammaproteobacteria, were decreased. At the genus level, *Ruminococcus, Eubacterium*, and *Butyrivibrio*, all classes of *Clostridia*, were increased (Table 2). Contrastingly, *Bacillus* and *Prevotella* were decreased after pretreatment. The increasing of Clostridia and decreasing other microbes caused the microbial diversity and richness decreasing (Table 3). Microbial diversity (Shannon index) was decreased to about 0.9 times while, microbial richness (OTU number and Chao1) more than 0.5 times decreased after pretreatment.

**Table 2.**
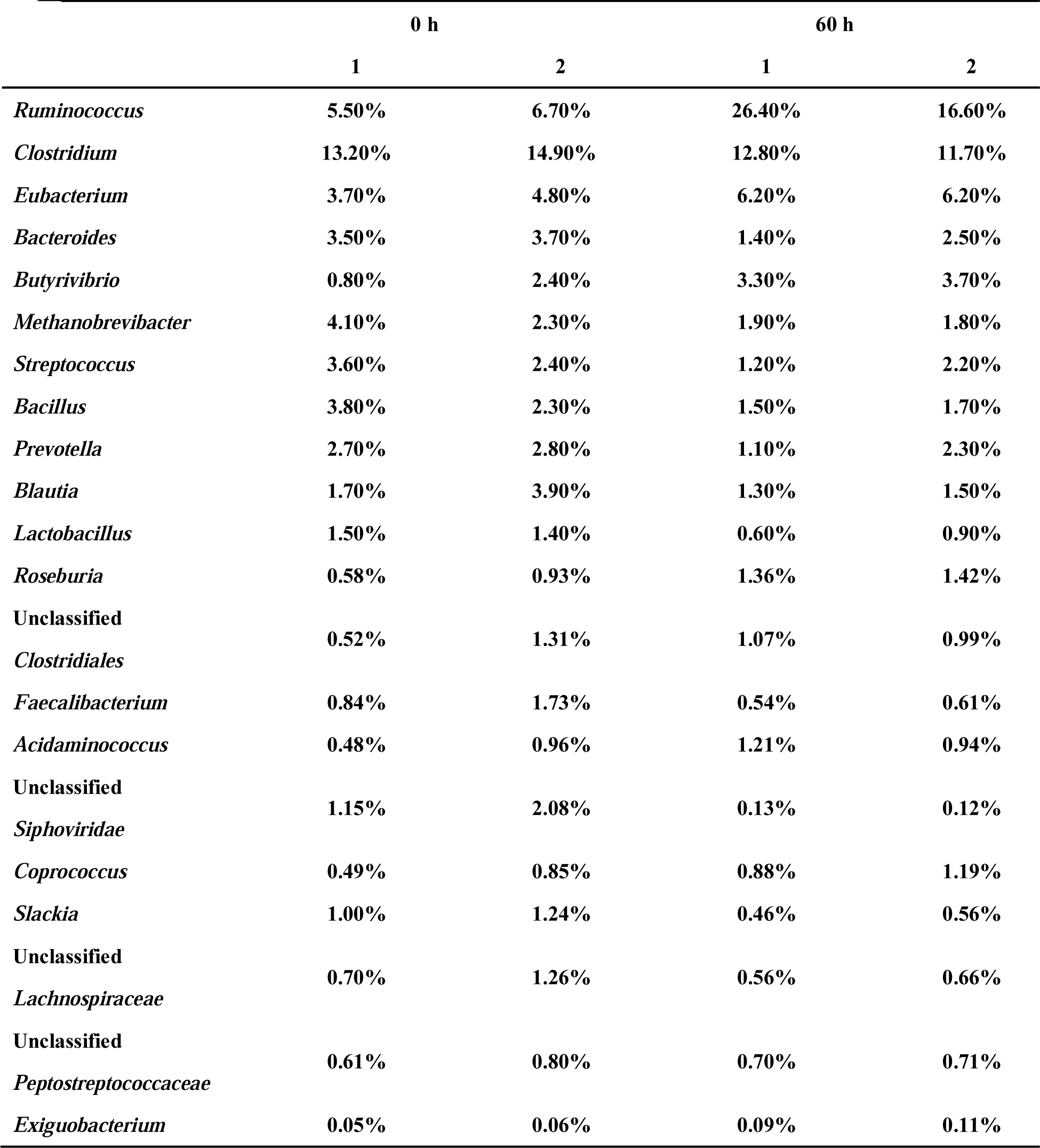
The relative abundance of top 20 genus and *Exiguobacterium* before and after pretreatment

**Table 3.** Microbial diversity and richness before and after pretreatment

|  | **0 h** |  | **60 h** |  |
| --- | --- | --- | --- | --- |
|  | **1** | **2** | **1** | **2** |
| **Shannon index** | **6.16** | **6.21** | **5.7** | **5.3** |
| **OTU number** | **2102** | **2223** | **2052** | **1011** |
| **Chao1** | **482** | **414** | **266** | **178** |

**Figure 2.**
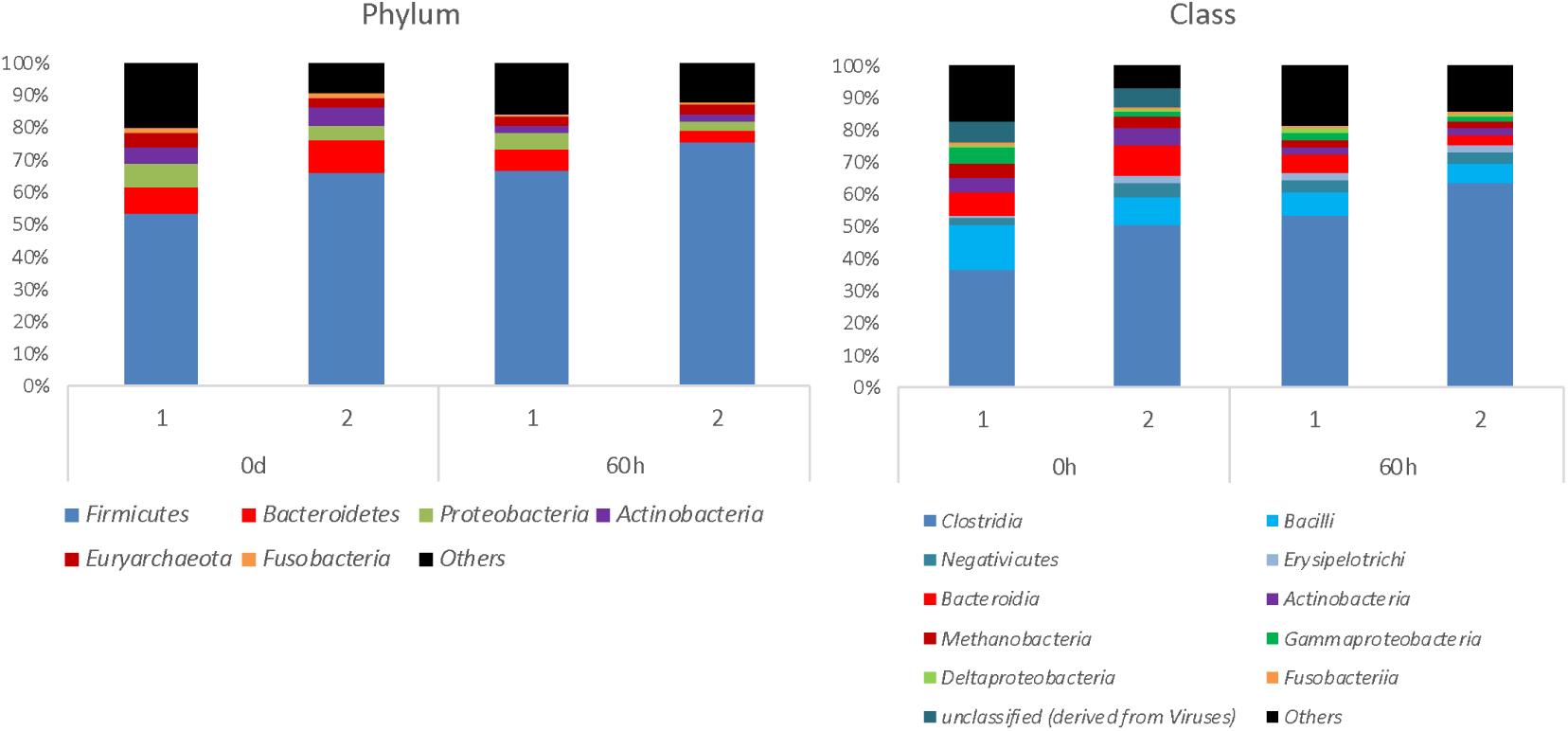
Relative abundance of microbial community (phylum and class level) during pretreatment.

### Prediction of gene functions and microbes involved in plant cell wall degradation

The functional binning of shotgun metagenomics (based on KEGG orthology) indicated that predominant functions detected in the rumen microbiota were metabolism (triphosphate reductase, phosphatase, etc.) at 56.8 % and genetic information processing (RNA polymerase, RNA synthetase etc.) at 25.5 %. Other functional assignments were linked to environmental information processes (11.8 %) and cellular processes (4.5 %).

We analyzed three genes encoding the proteins involved in cellulase and hemicellulase activities as beta-glucosidase (EC 3.2.1.21 and EC 3.2.1.74), beta-glucanase (EC 3.2.1.4), and endoxylanase (EC 3.2.1.8). Cellulose degradation (beta-glucosidase and beta-glucanase) collectively represented about 0.123 % of total sequence reads. The beta-glucosidase coding genes were mostly from bacterial genera *Butyrivibrio, Ruminococcus, Clostridium*, and *Exiguobacterium* (Figure 3). Interestingly, the relative abundance of *Exiguobacterium* was increased to about 100-fold and 40-fold in the sample after pretreatment, while that from *Bacillus, Butyrivibrio*, and *Prevotella* were decreased. Beta-glucanase was mainly affiliated with *Butyrivibrio, Clostridium*, and *Ruminococcus*. *Ruminococcus* genes were not detected in the rumen fluid prior to treatment. These genes increased while those from *Butyrivibrio* decreased after pretreatment. Endoxylanase was mainly assigned to *Clostridium* and *Ruminococcus* and both increased after pretreatment (Figure 4).

**Figure 3.**
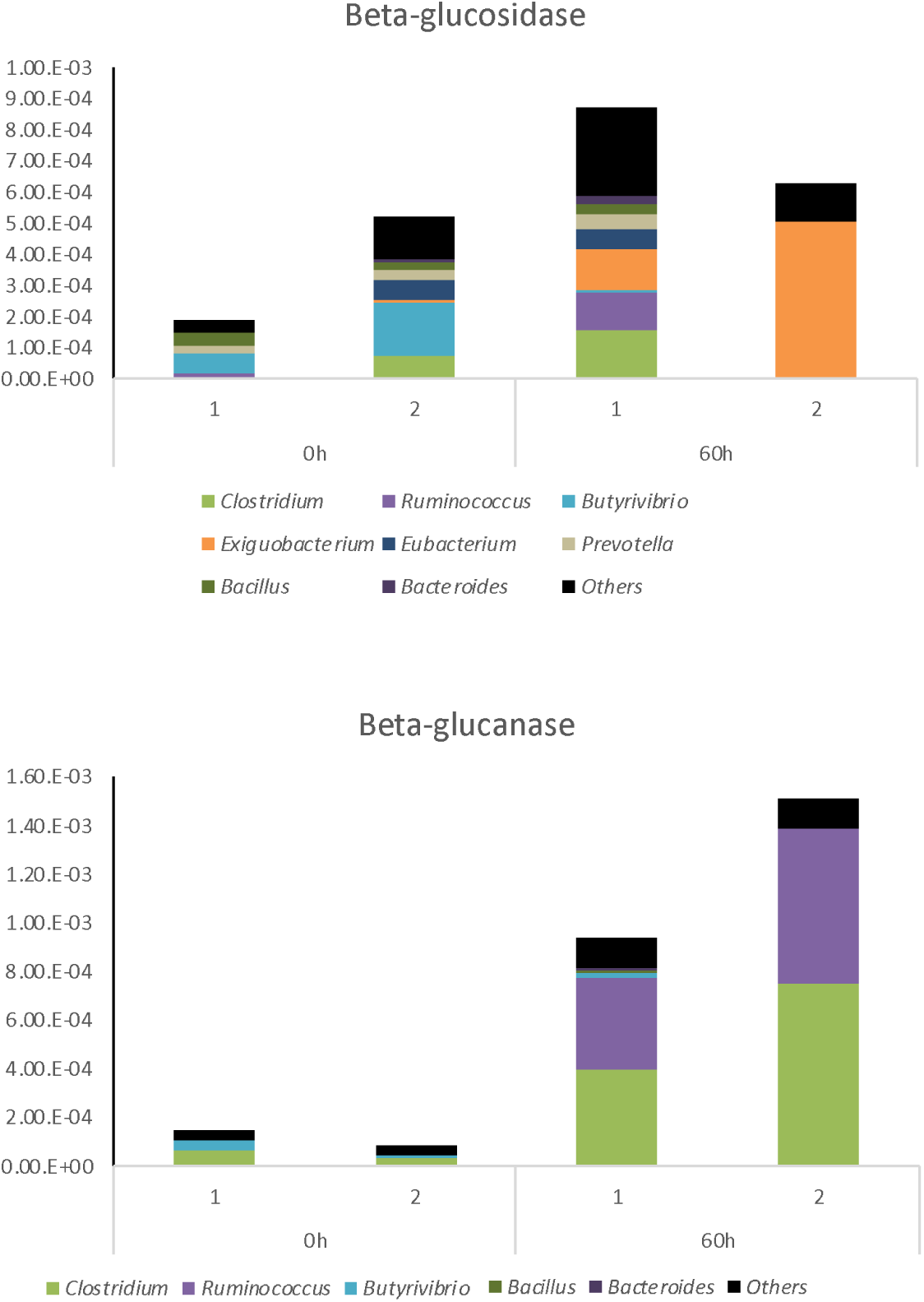
Relative abundance and taxonomic affiliation of cellulase (beta-glucosidase and beta-glucanase) during the incubation period.

**Figure 4.**
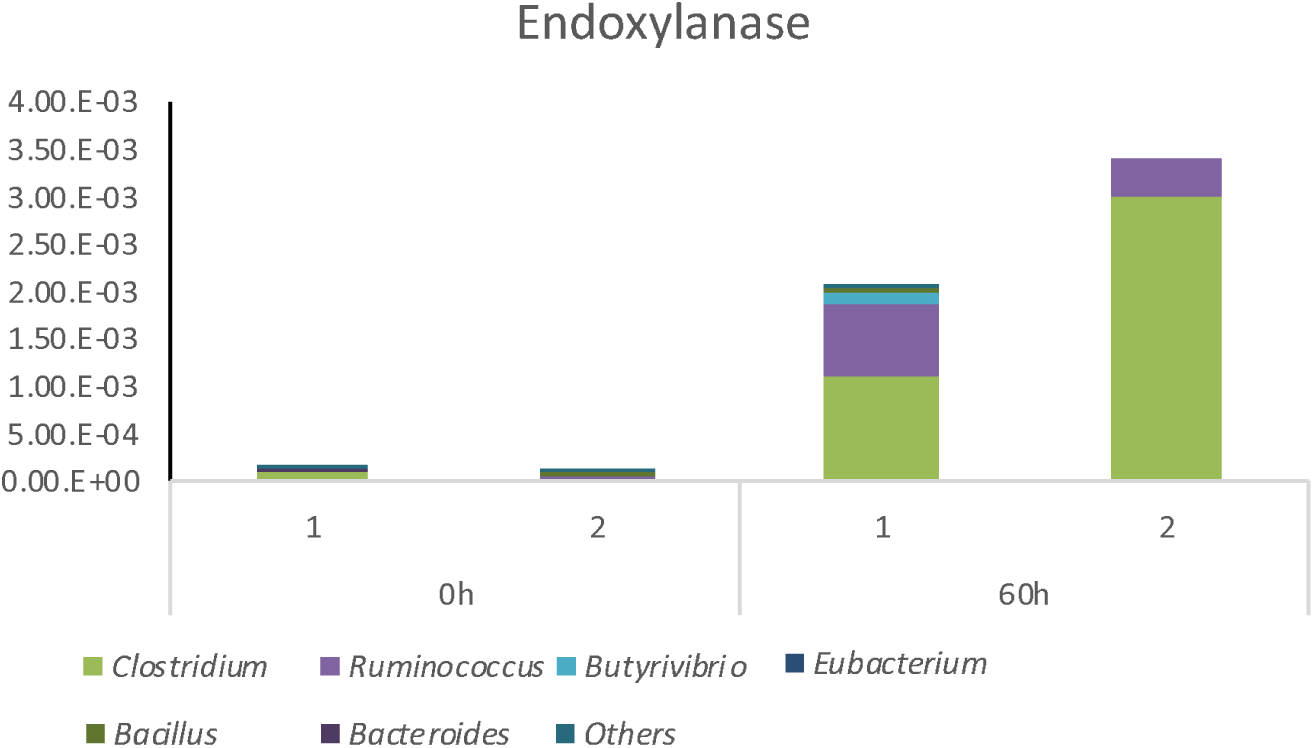
Relative abundance and taxonomic affiliation of hemicellulose (endoxylanase) during the incubation period.

## Discussion

In this study, we conducted metagenomic analysis to reveal the microbes involved in lignocellulose decomposition in the pretreatment system by cow rumen fluid. Culture-independent methods using a next-generation sequencer have been applied to study bacteria involved in plant cell wall degradation *in situ* cow rumen [3, 5, 11, 35]. In this study, we showed that *Ruminococcus* and *Clostridium*, they are belonging to *Clostridia*, were the primary cellulolytic and hemicellulolytic bacteria. Baba et al. [1] showed that after 24 h rapeseed (*Brassica napus* L.) pretreatment *Clostridia*, especially *Ruminococcus*, increased while *Bacteroidetes* decreased. Thus, *Clostridia* flourishing in the pretreatment system appear to be independent of substrate types. Microbial diversity and richness were decreased along with increasing of Clostridial abundance. *Clostridia* are gram-positive, anaerobic bacteria found in rumen that are capable of degrading plant cell walls [24, 37]. They are also known to be fibrolytic and several species have been isolated from rumen. *R. albus* and *R. flavefaciens*, are considered the major cellulolytic bacteria in the rumen [5, 10, 28, 36]. They also display degradation properties against of filter paper. In addition to total organism abundance, the cellulase and hemicellulase genes from *Clostridium* also increased after pretreatment. *C. thermocellum* and *C. aldrichii* can associated to the plant cell wall and have a cellulosome, which is a multienzyme complex specialized for cellulose degradation [18]. After the pretreatment, the beta-glucosidase genes affiliated with *Exiguobacterium* also increased.

*Exiguobacterium* has been detected with molecular biological methods but has not been isolated from rumen [5, 13]. They have been isolated from permafrost cores, soils, and several water environments, and they grow between −2.5 to 40 °C [26]. *Exiguobacterium* possess alpha-glucosidase, beta-glucosidase, and beta-galactosidase genes that enable them to decompose cellobiose and carboxymethyl cellulose [12, 26]. This is the first result showing the detection of the cellulase genes and their involvement in cellulose degradation from a cow rumen environment. *Clostridium* and *Exiguobacterium* are not dominant organisms (representing 5.0 and 0.01 % of *in situ* rumen flora, respectively) and not mainly involved in cellulose and hemicellulose degradation *in situ* rumen environment [3, 5]. However, they might mainly contribute to lignocellulose degradation in the pretreatment system despite they were not dominant. Thus, the detection of them as main lignocellulosic microbes *in vivo* rumen was unique finding compared with previous studies.

Plant cell wall degradation process *in situ* rumen is primarily attributed to *Ruminococcus, Prevotella, Butyrivibrio*, and *Fibrobacter* [3, 5, 19]. However, the genes encoding to cellulase and hemicellulase assigned to *Prevotella* and *Butyrivibrio*, were decreased after incubation. *Prevotella* belonging to *Bacteroidetes* are well known to be involved in decomposition of lignocellulosic biomass in a rumen [18, 30]. They can digest starch, hemicellulose, and pectin [4, 6]. *P. ruminicola* is a *Prevotella* strain found in rumen with endoxylanase and beta-glucosidase that can decompose filter paper [8, 20, 24]. *B. fibrisolvents* is also recognized as the main hemicellulolytic bacteria in the rumen [20]. Moreover, *P. ruminicola* and *B. fibrisolvents* can accelerate the cellulose digestion rate combined with fibrolytic bacteria such as *Ruminococcus* or *Fibrobacter* [17]. It has been reported that the quantities of *P. ruminicola* are increased significantly when the animal’s diet is switched from a hay to high grain, but that the abundance of *R. flavefaciens*, and *B. fibrisolvens* organisms are reduced [32]. Cellulose is the main components in waste paper, thus, non-cellulolytic *Prevotella* might be expulsed due to the dominance of cellulolytic *Ruminococcus, Clostridia*, and *Exiguobacterium*. The 16S rRNA based relative abundance of *Butyrivibrio* was increased after pretreatement, but functional genes-based cellulolytic *Butyrivibrio* was decreased. We were suggested that *Butyrivibrio* is not involved in cellulose or hemicellulose degradation, because *B. hungatei* or unknown *Butyrivibrio* spp. were increased.

In conclusion, we showed *Ruminococus* are primary members involved in plant cell wall decomposition, in agreement with previous studies. On the other hand, *Clostridium* and *Exiguobacterium*, which were primary fibrolytic bacteria in the previous studies, might play a more significant role in the decomposition of cellulose and hemicellulose in waste paper. Moreover, the main fibrolytic bacteria *in situ* rumen, *Prevotella* and *Butyrivibirio*, were also decreased. These results suggested the rumen pretreatment system environment was different to from the *in situ* rumen environment, and unique microbes were contributed degradation of lignocellulosic biomass. Future research will develop an effective decomposition system making use of *Ruminococcus, Clostridium*, and *Exiguobacterium.*

## Acknowledgment

We thank Mr. Kazuya Sato and Ms. Rie Yamamoto of Field Science Center, Tohoku University, for their technical support during collection of cow rumen contents. We would like to thank Editage (www.editage.jp) for English language editing. This work was supported by the Next-generation Energies for Tohoku Recovery (NET) project.

## Founding information

This work was supported by the Next-generation Energies for Tohoku Recovery (NET) project.

## Compliance with ethical standards

### Ethical approval

All experimental procedures conformed to “Regulations for Animal Experiments And Related Activities at Tohoku University", and were reviewed by the Institutional Laboratory Animal Care and Use Committee of Tohoku University, and finally approved by the President of University.

---

**Table S1.**
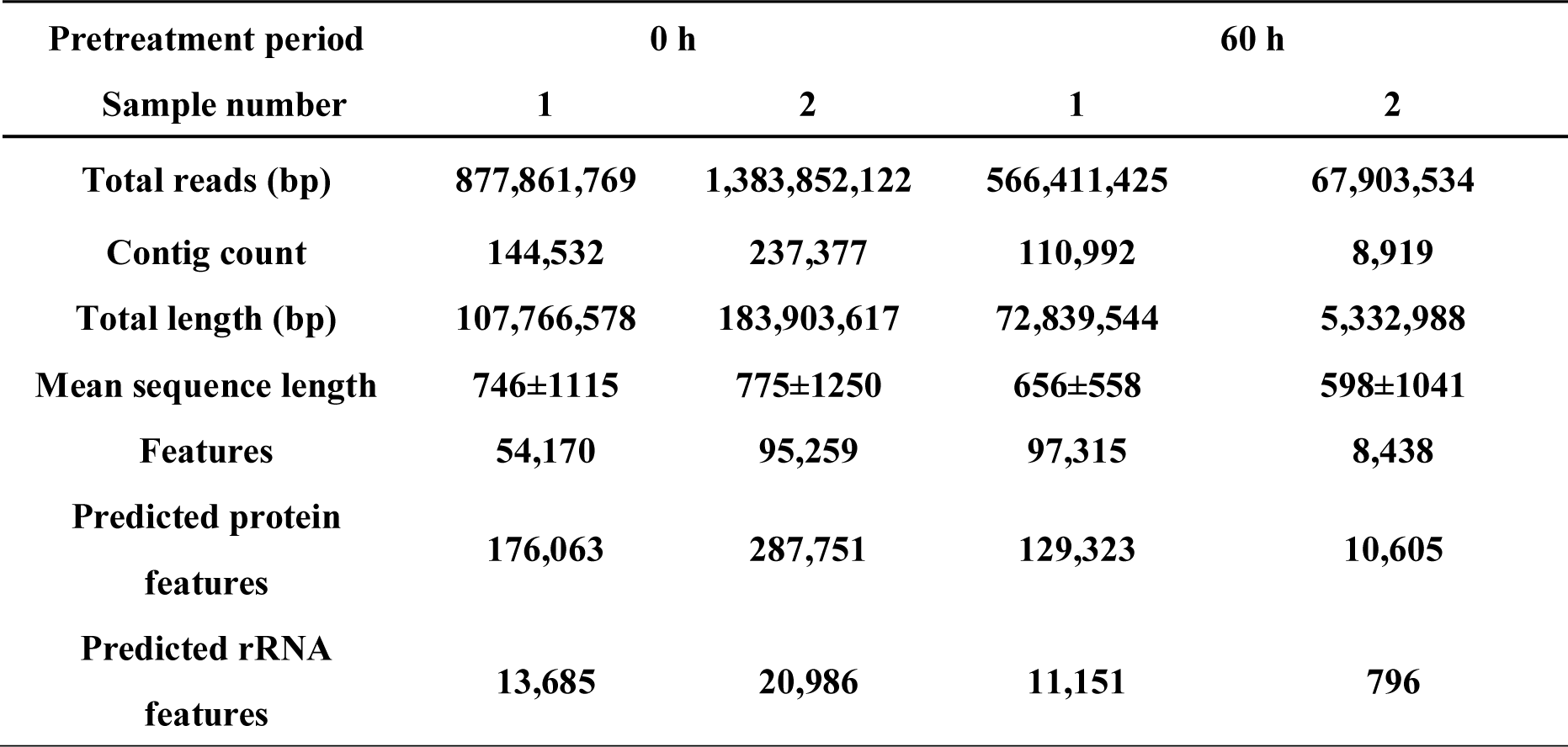
Results of shotgun metagenome sequencing analysis

